# Conservation of chromatin states and their association with transcription factors in land plants

**DOI:** 10.1101/2025.07.12.664529

**Authors:** Vikas Shukla, Elin Axelsson, Tetsuya Hisanaga, Jim Haseloff, Frédéric Berger, Facundo Romani

## Abstract

The complexity of varied modifications of chromatin composition is integrated in archetypal combinations called chromatin states that predict the local potential for transcription. The degree of conservation of chromatin states has not been established amongst plants, and how they interact with transcription factors is unknown. Here we identify and characterize chromatin states in the flowering plant *Arabidopsis thaliana* and the bryophyte *Marchantia polymorpha*, showing a large degree of functional conservation over more than 450 million years of land plant evolution. We used this new resource of conserved plant chromatin states to understand the influence of chromatin states on gene regulation. We established the preferential association of chromatin states with binding sites and activity of transcription factors. These associations define three main groups of transcription factors that bind upstream of the transcription start site, at the +1 nucleosome or further downstream of the transcription start site and broadly associate with distinct biological functions. The association with the +1 nucleosome defines a list of candidate pioneer factors we know little about in plants, compared to their important roles in animal stem cells and early development.

## INTRODUCTION

Eukaryotic genomes are organized into chromatin, a dynamic polymer of DNA and histone proteins that control access to the underlying genetic information. Chromatin structure, shaped by nucleosome positioning, histone modifications, histone variants, DNA methylation, and higher-order interactions, plays a fundamental role in gene regulation, genome stability, and chromosome organization. Advances in genome-wide profiling techniques have enabled the classification of chromatin states, defined by combinatorial patterns of histone marks and other epigenetic features, which reflect functionally distinct regulatory environments (1–3). Chromatin states provide a powerful framework to annotate genome function, delineate regulatory elements, and uncover specialised domains of chromatin within constitutive heterochromatin occupied by transposable elements (TEs), facultative heterochromatin occupied by genes transcriptionally repressed and euchromatin occupied by expressed genes.

A major function of chromatin is to regulate transcription by controlling the accessibility of DNA to transcription factors (TFs). TFs bind to specific cis-regulatory elements and influence gene expression by recruiting transcriptional machinery and chromatin-modifying complexes. In metazoans, models of chromatin states have proven especially useful in distinguishing functional modes of TF activity (4–6). Some TFs preferentially bind to open, transcriptionally active chromatin, while others engage nucleosomal DNA in repressive contexts, functioning as pioneer TFs that remodel chromatin to allow binding of additional TFs (6–9). These distinct modes of chromatin engagement are often predictive of TF mechanisms, such as the recruitment of the Polycomb repressive complex (PRC) (10–12) or histone acetyltransferases (HATs) (13,14).

In plants, chromatin state analysis has primarily focused on *Arabidopsis thaliana*, where recent efforts have defined high-resolution maps covering developmental stages and environmental conditions (15,16). This architecture shares analogous features with chromatin states in animals. These analyses constitute a valuable resource for functional genomics, but applications outside *Arabidopsis thaliana* are scarce (17,18) and the extent to which these combinatorial patterns of histone modifications are maintained across plant evolution remains an open question.

In contrast to animals, the interplay between TFs and chromatin has remained largely unexplored in plants. Early studies in *Arabidopsis thaliana* reported that most TFs bind open chromatin regions, with little variation in chromatin state preference across TF families (19), and the functional association between chromatin and TFs has remained unresolved. Notably, only a handful of TFs have been proposed to act as pioneer factors in plants. The best characterized so far is LEAFY (LFY), but other candidates among the MADS-box transcription factors have also been proposed as potential pioneer factors (20–22). Although hundreds of TF binding profiles have been generated in *Arabidopsis*, the field lacks a unified framework to relate these profiles to chromatin context and transcriptional activity. Genome-wide TF binding assays such as ChIP-seq or DAP-seq often reveal that many binding events are non-productive, i.e., they do not lead to measurable changes in gene expression (23,24). Unlike in animals, there is no unified chromatin–TF framework akin to ENCODE or modENCODE (25), complicating efforts to classify TFs based on binding context or regulatory output.

Here, by comparing *Arabidopsis thaliana* and *Marchantia polymorpha,* we showed the overall conservation of the role of histone H2A variants and histone post-translational modifications in defining conserved chromatin states. To characterize TF association with chromatin states, we defined a TF activity score aggregating both genome-wide TF binding data and co-regulation of gene expression. Using this strategy we showed that TFs can be grouped into distinct categories based on chromatin state preferences. These are involved in distinct biological functions, including a subset of candidate pioneer factors. We propose that TF–chromatin relationships follow similar principles in both species, supporting the conservation of regulatory strategies across land plants.

## METHODS

### Profiling of *M. polymorpha* H2A variants using ChIP-seq

ChIP experiments were performed using a previously described protocol with some modifications (26). Two-week-old thalli started from gemmae of *M*. *polymorpha* Tak- 1 wild type were collected and cross-linked using 1% formaldehyde in 1× PBS under vacuum on ice for 10 min. The cross-linking reaction was quenched by adding 2 M glycine to achieve a final concentration of 0.125 M under vacuum on ice for 10 min. Excess solution was removed from cross-linked tissue by blotting with paper towels. Cross-linked tissue was then snap frozen in liquid nitrogen and ground to a fine powder using mortar and pestle. The powder was transferred into a 50-ml plastic tube and suspended in 40 ml of MP1 buffer (10 mM MES-KOH pH 5.3, 10 mM NaCl, 10 mM KCl, 0.4 M sucrose, 2% (w/v) Polyvinyl pyrrolidone (PVP 10), 10 mM MgCl_2_, 10 mM 2-mercaptoethanol, 6 mM EGTA, 1× cOmplete protease inhibitor cocktail). Suspended samples were then filtered twice through one layer of Miracloth, once through a 40 μm nylon mesh, and twice through a 10 μm nylon mesh. Filtered samples were centrifuged at 3000 × ***g*** at 4°C for 10 min, and the supernatant was discarded. The pellet was washed using 15 ml of MP2 buffer (10 mM MES-KOH buffer pH 5.3, 10 mM NaCl, 10 mM KCl, 0.25 M sucrose, 10 mM MgCl_2_, 10 mM 2-mercaptoethanol, 0.2% Triton-X 100, 1× cOmplete protease inhibitor cocktail) three times. The final pellet was then resuspended in 5 ml of MP3 buffer (10 mM MES-KOH pH 5.3, 10 mM NaCl, 10 mM KCl, 1.7 M sucrose, 2 mM MgCl_2_, 10 mM 2-mercaptoethanol, 1× cOmplete protease inhibitor cocktail) and centrifuged at 16 000 × ***g*** at 4°C for 1 h. After centrifugation, the supernatant was discarded, and the pellet was resuspended in 900 μl of covaris buffer (0.1% SDS, 1 mM EDTA, 10 mM Tris–HCl pH 8.0, 1× cOmplete protease inhibitor cocktail). The resuspended pellet containing the chromatin fraction was fragmented using a Covaris E220 High-Performance Focused Ultrasonicator for 15 min at 4°C (duty factor, 5.0; peak incident power, 140.0; cycles per burst, 200) in a 1-ml Covaris milliTUBE. Sheared chromatin was centrifuged at 20 000 × ***g*** at 4°C for 10 min, and the supernatant was transferred into a new 5-ml tube and diluted by adding 2.7 ml of ChIP dilution buffer (0.01% SDS, 1.1% Triton-X-100, 1.2 mM EDTA, 16.7mM Tris-HCl pH 8.0, 167 mM NaCl). Diluted chromatin was cleared by incubating with proteinA/G beads (Thermo Fisher Scientific, Waltham, MA, USA) at 20 rpm rotating at 4°C for 1 h. Beads were removed by magnetic racks and precleared chromatin was separated into five tubes and incubated with 1 μg of specific antibodies for histone H2A variants at 20 rpm spinning at 4°C overnight. Chromatin bound by antibodies was collected by incubating with protein A/G beads for 3 h. The beads were collected by magnetic racks and washed twice with a low salt wash buffer (20 mM Tris–HCl pH 8.0, 150 mM NaCl, 2 mM EDTA, 1% Triton X- 100 and 0.1% SDS), once with a high salt wash buffer (20 mM Tris–HCl pH 8.0, 500 mM NaCl, 2 mM EDTA, 1% Triton X-100 and 0.1% SDS), once with a LiCl wash buffer (10 mM Tris–HCl pH 8.0, 1 mM EDTA, 0.25 M LiCl, 1% IGEPAL CA-630 and 0.1% sodium deoxycholate), and twice with a TE buffer (10 mM Tris–HCl pH 8.0 and 1 mM EDTA). Immunoprecipitated DNA was eluted using 500 μl elution buffer (1% SDS and 0.1 M NaHCO_3_) at 65°C for 15 min. To reverse cross-link, eluted DNA was mixed with 51 μl of reverse cross-link buffer (40 mM Tris–HCl pH 8.0, 0.2 M NaCl, 10 mM EDTA, 0.04 mg ml^−1^ proteinase K; Thermo Fisher Scientific) and incubated at 45°C for 3 h and then at 65°C for 16 h. After cross-link reversal, DNA was treated with 10 μg of RNase A (Thermo Fisher Scientific), incubated at room temperature for 30 min and purified using the MinElute PCR purification kit (Qiagen). ChIP-seq library was generated from ChIPed DNAs using Ovation® Ultralow Library Systems V2 (Tecan, Männedorf, Switzerland). The ChIP-seq libraries were sequenced on illumina Hiseq v4 to generate 50 bp single end reads.

### ChIP-seq data collection and processing

The publicly available ChIP-seq datasets for *Arabidopsis* were downloaded from our previous study (15). A detailed description of the sources of raw files can be found in the [source table 1]. The description of publicly available ChIP-seq datasets and the datasets generated for this study to calculate the chromatin states of *Marchantia* is also available in the [source table 1].

Raw BAM files were converted to FASTQ files using the “bamtofastq” sub- command of the “BEDTools suite” v2.27.1 (27). Sequencing quality of the raw files was evaluated using quality reports generated by FastQC v0.11.5 (28). Reads were trimmed using “Trim Galore” v0.6.5 (DOI:10.5281/zenodo.5127898). The trimmed reads were aligned to the reference genome: TAIR10 in case of *Arabidopsis* and MpTak-1 v7.1 in case of *Marchantia* using Bowtie2 v2.4.1 (29) and further processed using SAMtools v1.9 (30) and BEDTools v2.27.1 (27). Duplicates were removed using Picard v2.22.8 (https://broadinstitute.github.io/picard/) to generate the aligned BAM files. The correlation between ChIP samples was evaluated using the multiBamSummary and plotCorrelation functions of deepTools v3.1.2 (31). The bamCompare function of deepTools v3.1.2 was used to normalize the ChIP signal with Input/H3 and generate BigWig files with normalized ChIP signal. The code for the above-mentioned analysis can be found here: https://github.com/Gregor-Mendel-Institute/vikas_states_tf_study_2025.git.

### Chromatin state calculations

The aligned BAM files generated as explained above were used to calculate the chromatin states using the BinarizeBAM and LearnModel commands of ChromHMM v1.23 (2) with default parameters and models ranging from 2-50 states were learned. BAM files were binarized in 200 bp bins for model learning to characterize chromatin states using the multivariate Hidden Markov Model (HMM) to identify the combinatorial spatial patterns in the ChIP-seq signal of multiple chromatin marks. The models with 2 to 50 chromatin states were generated in each case. A 26-state model was chosen for the full *Arabidopsis* datasets based on two reasons: (1) As the raw data used to calculate states was the same as the one used in (15) with the main difference being the employment of stricter alignment parameters and filtering of previously included low confidence reads. These changes did not lead to extensive differences from the 26-state model in (15) [Supplementary Figure S1E]. A follow up on all models between 23 to 29 states revealed that models with less than 26 states showed loss of complexity (merging of states associated with distinct genomic features) and states with more than 26 states showed appearance of states that can no longer be associated with distinct genomic features. Hence, we decided to pick the 26-state model to be optimal for our analysis.

The chromatin states of *Marchantia* were calculated with the same pipeline. Given the comparatively smaller set of chromatin marks available in *Marchantia*, a smaller number of chromatin states was to be expected. A 15-state model was chosen for *Marchantia* using the similar strategy as in (15). To compare the chromatin state of *Arabidopsis* and *Marchantia*, we used only the chromatin marks that were available in both species and generated a less complex, comparable 15-state model for *Arabidopsis*.

### Annotation of chromatin states

The BED files of the chromatin states generated by ChromHMM were imported into R v4.3.2 (https://www.R-project.org/). The chromatin state regions were overlapped with the genomic features defined in *Arabidopsis* genome TAIR10 (GFF3 file downloaded from <u>here</u>) and *Marchantia* genome MpTak-1 v7.1 (GFF3 file downloaded from <u>here</u>) using the “bed_intersect” function of R package valr v0.6.6 (32). The results were plotted as stacked bar plots using the Tidyverse (33) package of R. [Figures 1B, 2B, and 3B]

**Figure 1.**
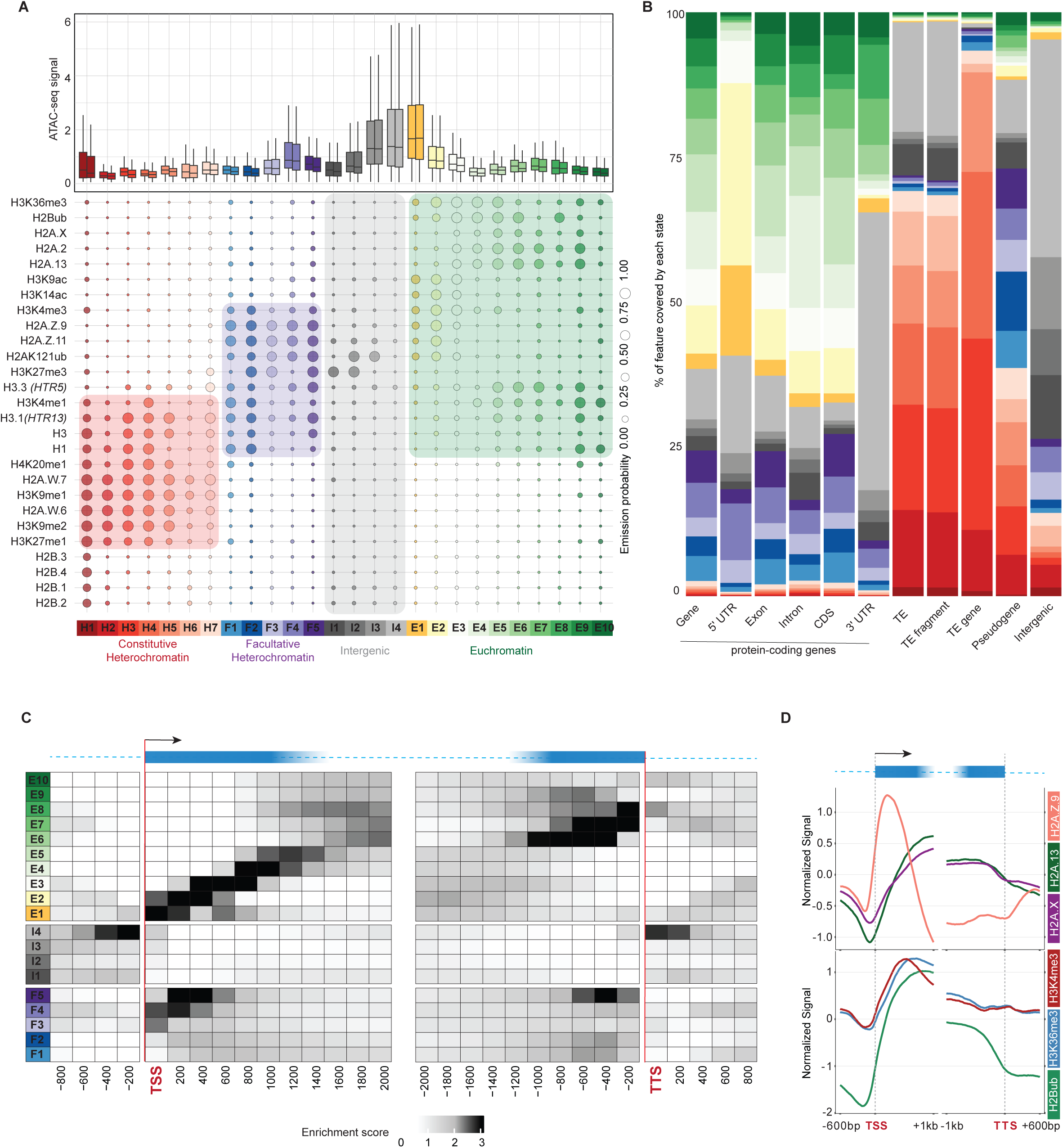
Chromatin states of *Arabidopsis thaliana*. **(A)** Bubble plot showing the emission probabilities for histone modifications/variants across the 26 chromatin states of *Arabidopsis thaliana*. The size of the bubble represents the emission probability ranging from 0 to 1. Coloured rectangles demarcate the classification of states into major domains of chromatin. The box plot on top shows the average ATAC-seq signal for each state representing chromatin accessibility. The two boxes per state are the two replicates of the ATAC-seq experiment. **(B)** Stacked bar plot showing the overlap between annotated genomic features and chromatin states. **(C)** Neighbourhood Enrichment analysis of chromatin states representing the fold enrichment for each state at fixed positions relative to the anchor position (TSS and TTS). **(D)** Metaplot showing the enrichment of prominent histone variants and modifications associated with active transcription. The enrichments here are plotted on genes overlapping states E1-E8. (n = 15234 / 27416 protein coding genes)

The NeighbourhoodEnrichment command of ChromHMM v1.23 (2) was used to calculate the enrichment of each state relative to the Transcription Start Site (TSS) as anchors that were extracted from aforementioned GFF files. The resultant matrix was imported into R v4.3.2 (https://www.R-project.org/) and the heatmap was plotted using the Tidyverse (33) package of R. [Supplementary Figures S1A, S2A, and S3A]

To evaluate the genomic localization of states, especially the heterochromatic states (H1:H7) in *Arabidopsis*, histograms with binwidth of 50,000 bp were plotted on chromosome 1 using the Tidyverse (33) package of R showing the preferential localization of heterochromatic states near the centromere region. [Supplementary Figures S1F and S2F]

We identified the states of constitutive heterochromatin by their strong enrichment in the regions containing Transposable Elements (TE), TE genes and TE fragments [Figures 1B, 2B, and 3B] along with high emission probabilities of know heterochromatic marks including H2A.W, H3K9me1/2 among others. [Figures 1A, 2A, and 3A] The distinction between the states of euchromatin and facultative heterochromatin was made by the high emission probabilities of marks associated with each type of chromatin state. The states of intergenic region showed little to no enrichment of any specific mark, representing Nucleosome Free Regions (NFR) and constituted about 20% of the genomes in both *Arabidopsis* and *Marchantia*. [Supplementary Figures S1B, S2C, and S3B]. We decided to annotate the states with the following nomenclature strategy:

(1) Euchromatin: The states of euchromatin were named with letter “E” in *Arabidopsis* 26-state model, with letter “e” in *Arabidopsis* 15-state model, and with letters “MpE” followed by a number in *Marchantia* 15-state model. The order of the numbers was assigned based on their enrichment relative to the TSS going from TSS to gene body. [Supplementary Figures S1A, S2A, and S3A]
(2) Constitutive Heterochromatin: The states of constitutive heterochromatin were named with letter “H” in *Arabidopsis* 26-state model, with letter “h” in *Arabidopsis* 15-state model, and with letters “MpH” in *Marchantia* 15-state model. The order of the numbers was assigned based on their localization on chromosomes going from centromere to peri-centromere in case of *Arabidopsis* states. [Supplementary Figures S1F and S2F]
(3) Facultative Heterochromatin: The states of facultative heterochromatin were named with letter “F” in *Arabidopsis* 26-state model, with letter “f” in *Arabidopsis* 15-state model, and with letters “MpF” in *Marchantia* 15-state model.
(4) Intergenic states: The states of intergenic regions were named with letter “I” in *Arabidopsis* 26-state model, with letter “i” in *Arabidopsis* 15-state model, and with letters “MpI” in *Marchantia* 15-state model.

### Analysis of chromatin states

The emission matrices generated by ChromHMM were imported into R v4.3.2 (https://www.R-project.org/). The state names and colours were reassigned using the strategy aforementioned, followed by plotting the emission matrices as bubble plots using the Tidyverse (33) package of R. [Figures 1A, 2A, and Supplementary Figure S3A]

The percentage of genome covered by each state was calculated as follows: (Total number of base pairs covered by each state / Genome size) * 100. The results were plotted as a pie chart using the Tidyverse (33) package of R. [Supplementary Figures S1G, S2A, and S3C]

We compared the chromatin states published in (15) with our 26-state model by importing the BED files for both models into R followed by overlapping the two matrices to create the transition matrix containing the occurrences of state transition between the two models. The data was plotted as an alluvial plot using the “circlize” (34) package of R. Similar analysis also led to the comparison of the *Arabidopsis* 26 and 15-state models. [Supplementary Figures S1E and S3I]

To show the similarities between the states of *Arabidopsis* 15-state model and the states from extensive 26-state model, after removing the additional marks from the 26- state model that were missing in the 15-state model.

To show the similar association of chromatin marks between the states of *Arabidopsis* and *Marchantia* 15-states models, hierarchical clustering of the two emission matrices was performed. H2A.W of *Arabidopsis* and H2A.M.2 of *Marchantia* were used interchangeably for comparison purposes. The two matrices were imported into R and merged for the common marks. The heatmap and clustering was performed using ComplexHeatmap (35) package of R. [Supplementary Figure S2G]

### RNA-seq, ATAC-seq, and Bisulfite-seq data analysis

The Bisulfite-seq and ATAC-seq data for wild type *Arabidopsis* seedlings was downloaded from GEO accession: <u>GSE146948</u> (36) while the RNA-seq data was downloaded from (37). All data was aligned to TAIR10 using bowtie2 v2.4.1 (29) and further processed using SAMtools v1.9 (30) and BEDTools v2.27.1(27). BigWig files for ATAC-seq and Bisulfite-seq and counts file for RNA-seq was imported into R. The data was overlapped with states regions file using the valr v0.6.6 (32). The results were plotted as box plots using the Tidyverse (33) package of R [Figure 1A, and Supplementary Figures S1A-D, S3D-G].

The Bisulfite-seq, ATAC-seq, and RNA-seq data for *Marchantia* was downloaded from (38) accession numbers SRA: PRJNA553138 and PRJDB8530. All data was aligned to MpTak-1 v7.1 genome and the boxplots were generated with the pipeline mentioned above [Figure 2A, and Supplementary Figures S2B-E].

**Figure 2.**
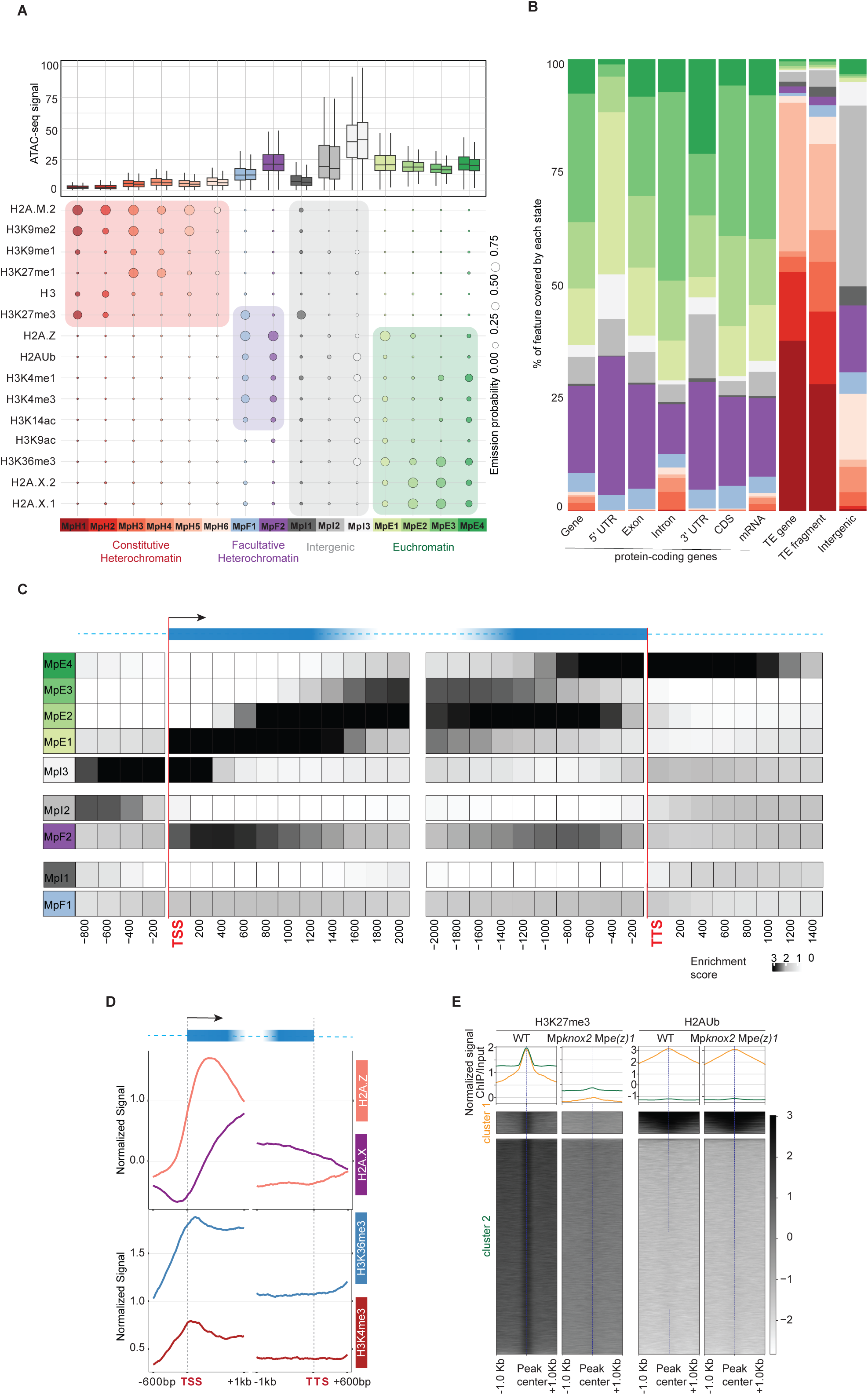
Chromatin states of *Marchantia polymorpha*. **(A)** Bubble plot showing the emission probabilities for histone modifications/variants across the 15 chromatin states of *Marchantia polymorpha*. The size of the bubble represents the emission probability ranging from 0 to 1. Coloured rectangles demarcate the classification of states into major domains of chromatin. The box plot on top shows the average ATAC-seq signal for each state representing chromatin accessibility. The two boxes per state are the two replicates of the ATAC-seq experiment. **(B)** Stacked bar plot showing the overlap between annotated genomic features and chromatin states. **(C)** Neighbourhood Enrichment analysis of chromatin states representing the fold enrichment for each state at fixed positions relative to the anchor position (TSS and TTS). **(D)** Metaplot showing the enrichment of prominent histone variants and modifications associated with active transcription. The enrichments here are plotted on genes overlapping states MpE1-MpE4. (n = 10715 / 17944 protein coding genes) **(E)** Enrichment heatmap and metaplots showing the apparent disassociation between H3K27me3 and H2AKUb marks in wild type and Mp*knox2* Mp*e(z)1* double mutants. The enrichment of both marks is plotted on wild type H3K27me3 peaks (n = 25159). The peaks have been clustered using k-means clustering into two clusters where cluster 1 represents peaks of H3K27me3 shared with H2AKUb while cluster 2 represents majority of H3K27me3 peaks that do not overlap with H2AKUb.

### Association of chromatin states with expression

To assess the relationship between chromatin states and gene expression, we extracted promoter sequences from *Arabidopsis* thaliana (TAIR10) and *Marchantia* polymorpha (Tak1 v7.1), defining promoters as the 1,000 bp region upstream of the transcription start site (TSS) using the promoter function from the GenomicRanges package (v1.56) in R. For each gene in *Marchantia* genome, we determined the overlap of these promoter regions with annotated chromatin states using the findOverlaps and pintersect functions.

For gene expression data, we compiled a diverse set of publicly available RNA-seq experiments from MarpolBase expression (39) for *Marchantia* and EvoRepro for *Arabidopsis* (40). To examine chromatin state association with transcriptional output, we ranked genes by their expression levels—measured in transcripts per million (TPM)—in the primary vegetative tissues where chromatin state data were derived (*Arabidopsis* leaf and *Marchantia* thallus). Genes were grouped into 20 equal-sized bins based on TPM values, and for each bin, we calculated the average proportion of each chromatin state per bin and visualized the distribution.

Similarly, to assess chromatin state dynamics across broader expression datasets, we analysed the same set of RNA-seq samples and estimated transcriptional variability by calculating the coefficient of variation (CV) for each gene, defined as the ratio of the standard deviation to the mean TPM across samples. Genes were ranked by CV, divided into bins, and plotted as described above.

For tissue-specific expression, we identified genes with at least a two-fold change in expression relative to their average expression across all tissues and assessed chromatin state distributions accordingly. A similar approach was applied to analyse chromatin state occupancy over gene bodies. To identify genes marked by H3K27me3, we applied the method described in (38) with data from (38) and (41).

### Transcription Factor Occupancy and Chromatin State Association

To analyse TF occupancy, we compiled publicly available *Arabidopsis* thaliana ChIP-seq and DAP-seq BED files from multiple studies (19,42,43). For *Marchantia* polymorpha, we inferred TF binding sites by mapping position weight matrices (PWMs) from (42) to the *Marchantia* v7.1 genome using FIMO (MEME Suite) (44). In both cases, each significant peak region in the genome was classified according to chromatin state using the OverlapsAny function from GenomicRanges package in R. For that purpose, only the centre of the peak was used to assign the chromatin state.

TF binding enrichment for each chromatin state was calculated following (6). Specifically, for each TF and chromatin state (s), enrichment was determined using: (a_s_/b)/(c_s_/d), where a_s_ is the total number of bases in a peak call in s; b is the total number of bases in a peak call; c_s_ is the total number of bases in s; d is the total number of bases of the chromatin state.

To calculate TF activity, we incorporated gene co-expression data. First, genes associated with TF peaks were annotated using the ChIPseeker package (45) with default parameters. Then, co-expression data from ATTEDII (46) (version Ath-r.c3-1) was used to filter TF target genes. Both positively (score < 2000) and negatively (score > max(score) - 1000) co-expressed genes were retained. The same enrichment formula described above was applied to this filtered subset. After this filter, we used the same formula than before.

To calculate the score of TF activity, we assessed statistical Enrichment of co-Expressed TF targets in chromatin states, we performed Fisher’s exact tests (alternative hypothesis = greater). For each chromatin state, we compared TF targets with positively or negatively co-expressed genes as described above. Peaks were annotated as described above, and gene IDs were extracted for each chromatin state. As null distribution, we considered a random sample of genes from the whole genome using this contingency table:

|  |  |
| --- | --- |
| Count (genes with peaks + co<br>expressed) | count (genes with peaks) |
| Count (co expressed) | 27205 for *Arabidopsis* or<br>18328 for *Marchantia* |

The TF-chromatin state score was computed as: -log_10_(p-value) + log_2_(odds ratio + 1) and normalized using the maximum score for each TF.

For each method we generated an N_TF_ × N_state_ matrix containing all computed scores and visualized chromatin state occupancy patterns using the pheatmap R package. For aggregate analyses across all TFs, scores were summed and normalized to the maximum score for each state.

In subsequent analyses, TFs with maximum scores below 8 were filtered out to exclude low-confidence results. TF family enrichment was assessed using Fisher’s exact tests, employing TF classifications from (42) and PlantTFDB (47). GO term enrichment analysis of biological processes in *Arabidopsis* was performed using PANTHER (48) with default parameters.

## RESULTS

### Chromatin states exhibit positional preference for gene regulatory regions of *Arabidopsis thaliana*

Chromatin states integrate the specific combinatorial occupancy of histone modifications and variants (2) at a near nucleosome resolution (bin size = 200 bp; see Methods). These states help delineate major functional domains of chromatin and provide insights into transcriptional regulation. We independently used ChromHMM (2) to recalculate the chromatin states of the *Arabidopsis thaliana* genome (TAIR 10) using the series of genomic profiles of histone variants and most abundant histone PTMs used by (15). We fine-tuned the chromatin state annotation pipeline and used stricter quality control steps (see Methods), to obtain a refined and more robust definition of chromatin states (15,16). Each group of states were defined by a specific combination of histone variants and histone PTMs [Figure 1A] and were associated with diverse levels of transcripts [Supplementary Figure S1A], DNA methylation [Supplementary Figures S1B – D], and were differentially associated with transposable elements, genes or intergenic regions [Figure 1B]. The new model contained 26 states including seven states of constitutive heterochromatin (H1 – H7) on transposable elements, four intergenic states (I1 – I4), five states of facultative heterochromatin (F1 – F5) on lowly expressed protein coding genes, and ten states in euchromatin on expressed genes (E1 – E10) [Figure 1A]. The enrichment in specific classes of H2A variants marked the identity of each group of states, with H2A.W in constitutive heterochromatin, H2A.Z in facultative heterochromatin, and H2A (and H2A.X) on expressed genes. Only minor differences separated this set of chromatin states from those previously described (15). The previous states H4 becomes annotated as H2, and the previous states H2 and H3 split in states H3 – H5, and the reannotation of the facultative chromatin states including the state F1 split into I1 and I2 [Supplementary Figure S1E].

We investigated the positional preference of chromatin states along chromosomes and observed that the states of constitutive heterochromatin (H1-H7) were primarily enriched on transposable elements and showed a positional preference with respect to the centromere of chromosomes [Supplementary Figure S1F]. The state H1 showed an exclusive preference for centromeric chromatin while states H2 to H7 were successively enriched from pericentromeres (H2-H5) to chromosome arms (H6 and H7) [Supplementary Figure S1F]. This constitutive heterochromatin represents ca. 20% of the genome [Supplementary Figure S1G].

The other states were primarily enriched at genes, and we performed neighbourhood enrichment analysis with the Transcription Start Site (TSS) or the Transcription Termination Site (TTS) as anchors. [Figure 1C; see Methods]. In euchromatin, expressed genes were occupied by a remarkable sequential enrichment pattern of chromatin states. The state I4 was most specifically enriched in the promoter and terminator regions of most protein-coding genes while the states E1 - E8 were organized from the promoter towards the gene body. Each nucleosome occupies ca. 200 bp and the states E1, E2, and E3 showed enrichment in positions corresponding to the 1^st^, 2^nd^, and 3^rd^ nucleosomes, respectively, for most of the expressed protein-coding genes. The states E4 and E5 occupied gene bodies and E5 was also marked by high CG methylation and enrichment in H3.3 compared with state E4 [Figures 1A, 1C and Supplementary Figure S1B], states E6, E7, and E8 occupied the end of the gene body [Figure 1C]. Ploting the enrichment of major histone H2A variants and chromatin modifications over genes occupied by states E1-E8 supported the predominance of H2A.Z associated with H3K4me3 in the first nucleosomes while the gene body is occupied by H2A and H2A.X with H3K36me3 and H2bUb [Figure 1D]. A succession of facultative heterochromatin states F3 – F5 occupied gene bodies [Figure 1C] although their organization was less complex compared with euchromatin states. The facultative heterochromatin states were primarily associated with the intergenic states I1 to I3, based on their enrichment in H3K27me3 and H2AK121ub, low accessibility, and low gene expression [Figure 1A and Supplementary Figure S1A] and altogether occupied circa 30% of the genome [Supplementary Figure S1G]. Contrasting with these graded patterns of chromatin states, the states E9 and E10 (ca. 5% of euchromatin) and the states F1 and F2 did not show a strong positional enrichment relative to TSS or TES (Fig 1C), but shared enrichment in linker histone H1 and were associated with genes expressed at very low levels [Figures 1A, 1C and Supplementary Figure S1A], suggesting a specific role of H1 in defining these chromatin states in addition to the major role of histone PTMs and H2A variants.

## Conservation of chromatin states in land plants

To investigate the conservation of chromatin states in land plants, we used the same approach as above to define chromatin states in *Marchantia polymorpha*. This model liverwort shares a similar genome organization with model species of hornworts and mosses that, together, form the monophyletic group of bryophytes, which diverged from vascular plants about 450 mya (49). To the set of publicly available histone-modification enrichment profiles (38,50), we added profiles of histone H2A variants detected with specific antibodies for H2A.X.1, H2A.X.2, H2A.Z, and H2A.M.2, which are expressed in vegetative tissues of *Marchantia* (26). Enrichment of chromatin features was mapped onto the newest telomere-to-telomere chromosome-level genome assembly v7.1 (51). Using ChromHMM we described fifteen chromatin states [Figure 2A; see Methods] which were annotated based on their differential enrichment in different genomic features [Figure 2B]. Intergenic states covered approximately a quarter of the genome [Supplementary Figure 2A]. Chromatin states associated with specific levels of gene expression [Supplementary Figure S2B], chromatin accessibility [Figure 2A], and DNA methylation [Supplementary Figure S2C - E]. Based on the enrichment of known constitutive heterochromatin marks, including H3K9me1/2 and H3K27me1, we ascribed six distinct states to constitutive heterochromatin (MpH1 - MpH6) [Figure 2A] that covered 39% of the genome [Supplementary Figure S2A]. These states were also distinguished by the lowest accessibility, enrichment in the H2A variant H2A.M.2 [Figure 2A], the highest degree of DNA methylation [Figures S2C - E] and were primarily present on transposable elements, with a minor representation on protein coding genes [Figure 2B]. The hallmark of heterochromatin states is their enrichment in H2A.M.2 and H3K9me2, but they are divided into three groups [Figure 2A]. MpH5 and MpH6 do not carry additional marks, except for methylated cytosines in MpH5, while states MpH3 and MpH4 are enriched in H3K9me1 and H3K27me1. Unlike heterochromatin states in *Arabidopsis*, *Marchantia* heterochromatin states MpH1 and MpH2, which represent more than a third of constitutive heterochromatin are also enriched in H3K27me3 with less DNA methylation than MpH3 and MpH4.

Facultative heterochromatin states MpF1 and MpF2 both enriched in H2A.Z and H2A ubiquitination (H2AUb) were associated with the lowest range of gene expression covered 16 % of the genome [Figures 2A, 2B and Supplementary Figures S2A, and S2B] and the state MpF1 also enriched in H3K27me3. As in *Arabidopsis*, facultative heterochromatin states were also enriched in H3K4me1 and H3K4me3. The neighbourhood analyses centred on TSS and TTS showed that the association of intergenic states MpI1 upstream of MpF1, both enriched in H3K27me3, representing one third of facultative heterochromatin with low accessibility occupying primarily repressed genes [Figures 2A-C and Supplementary Figures S2A, S2B]. The remaining two thirds of facultative heterochromatin of *Marchantia* was occupied by the states MpI2 and MpF2, marked by an enrichment of H2A.Z and H2AUb but not H3K27me3 and occupying more accessible chromatin and genes less repressed than genes covered by F1 [Figures 2A, 2C, 2D and Supplementary Figure S2B]. MpI2 occupies 19% of the genome and altogether facultative heterochromatin occupies 37% of the genome which is much higher than in *Arabidopsis*.

Regions with the highest gene expression represented euchromatin [Supplementary Figure S2A] and comprised the intergenic state MpI3 and four genic euchromatin states E1 – E4 covering altogether with 24 % of *Marchantia* genome [Figures 2A, 2B and Supplementary Figure S2A]. The neighbourhood analyses centred on TSS and TTS showed that the intergenic state MpI3 corresponds to a region located in intergenic regions upstream of the TSS [Figures 2B, 2C]. Contrary to *Arabidopsis,* the promoters of *Marchantia* defined by the region just upstream of the TSS showed enrichment of H2AUb and the elongation mark H3K36me3, along with other euchromatic marks. The most accessible, non-intergenic state is MpE1, which marked the +1 nucleosome of most protein-coding genes of *Marchantia* [Figures 2A, 2B]. This state, with strong enrichment of H2A.Z and H3K36me3, is present in the first few nucleosomes of active genes while the states MpE2 to MpE4 successively occupy the gene body and are enriched in H2A.X [Figures 2C, 2D]. To evaluate the relative role of histone H2A variants and histone PTMs in defining chromatin states, we calculated a new matrix of chromatin states in the absence of data on either H2A variants or histone PTMs and used the Jaccard index to compare both resulting matrices to the original 15 chromatin states [Supplementary Figure S2F]. A high Jaccard index reports high similarity. We observed that *in silico* removal of H2A variants perturbed the definition of constitutive heterochromatin states less strongly than removal of histone PTMs while the influence of H2A variants and histone PTMs was comparable on the chromatin states of facultative heterochromatin and euchromatin.

To directly compare the chromatin states of *Arabidopsis* and *Marchantia*, we first calculated a 15-states model in *Arabidopsis* based on the same set of histone modifications and H2A variants used to identify chromatin states of *Marchantia*. The 15 and the 26 states models were highly comparable with a notable reduction of the complexity of constitutive heterochromatin to only four states and of euchromatin to only five states [Figure 1A, Supplementary Figure S3A and S3I]. To compare the 15 chromatin states of *Arabidopsis* and *Marchantia*, we performed hierarchical clustering of emission probabilities of the two models [Supplementary Figure S2G]. The resulting tree supported the overall similarities of the constitutive heterochromatin and euchromatin states. In contrast, while the typical H3K27me3 enriched in facultative heterochromatin states of the two species clustered, the *Marchantia* facultative heterochromatin MpF2 clustered with the *Arabidopsis* euchromatic states E2, which was likely due to the prominent enrichment in H2A.Z in their definition. We noted that chromatin accessibility of MpI2 and MpF2 were more similar to accessibility of euchromatin states than in *Arabidopsis* [Figure 2A, Supplementary Figure S3A] suggesting an ambivalent role for this subtype of facultative heterochromatin in *Marchantia*. To further compare the regulation by PRC2 in both species, we analysed chromatin landscapes in the *Marchantia* Mp*knox2* Mp*e(z)1 double* mutant, which exhibited reduced PRC2 activity (41,50). Peaks enriched in H3K27me3 in the wild type were depleted in the mutant deprived of PRC2, but the loss of PRC2 activity in *Marchantia* did not affect the deposition of H2AUb [Figure 2E] presumably by PRC1 (52). These findings suggest a strong independence of the two Polycomb repressive pathways in *Marchantia*.

While our data supported an overall conservation of the chromatin states between the two species, we uncovered unique features of *Marchantia* chromatin organization. The bulk of the modification H3K27me3 was associated with transposable elements and only to a minor fraction of facultative heterochromatin. Instead, H2A.Z and H2AKUb were the dominant marks of facultative heterochromatin, which is consistent with the mild effect of the loss of H3K27me3 on vegetative development (50) compared with *Arabidopsis* (53,54). The promoter state MpE1 enriched in the transcription elongation mark H3K36me3, and the long persistence of E4 after the TTS creates an asymmetry between the intergenic regions upstream and downstream of each gene. This is in contrast with *Arabidopsis*, where the intergenic state I4 symmetrically occupies these two regions [Figure 2C]. This further suggests that in *Marchantia*, the orientation of genes defines distinct chromatin environment in their vicinity, through mechanisms yet to be uncovered. The distinct positional preferences of chromatin states in *Arabidopsis* versus *Marchantia*, particularly their sequential organization around gene regulatory regions, suggested a functional relationship with transcriptional regulation. To examine this in further detail, we ranked genes by their expression levels in *Arabidopsis* seedlings and *Marchantia* thalli. Highly expressed genes were depleted of facultative chromatin states [Figure 3A]. In the case of *Arabidopsis*, I3 was also depleted in highly expressed genes and enriched in genes with low expression [Figures 3A and 3D]. The E1 state, which was close to the TSS, was enriched in highly expressed genes in both species but more prominently in *Marchantia*, where it included the proximal promoter [Figures 3A and 3D]. Genes with low expression in vegetative tissues may be transcriptionally active at specific developmental stages or under environmental stimuli. To explore this, we assessed the coefficient of variation (CV) of gene expression across a panel of RNA-seq datasets spanning *Marchantia* and *Arabidopsis* development. The CV, defined as the standard deviation divided by the mean expression level, is independent of absolute expression and distinguishes ubiquitously expressed genes from those with variable expression in different conditions. Genes with high CVs exhibited greater coverage of facultative chromatin states, whereas ubiquitously expressed genes were largely depleted of these states [Figures 3B and 3E]. In contrast, euchromatin states showed the opposite trend. A similar pattern was observed when analyzing chromatin states within coding regions, with a much more prominent participation of euchromatin states [Supplementary Figure S4].

**Figure 3.**
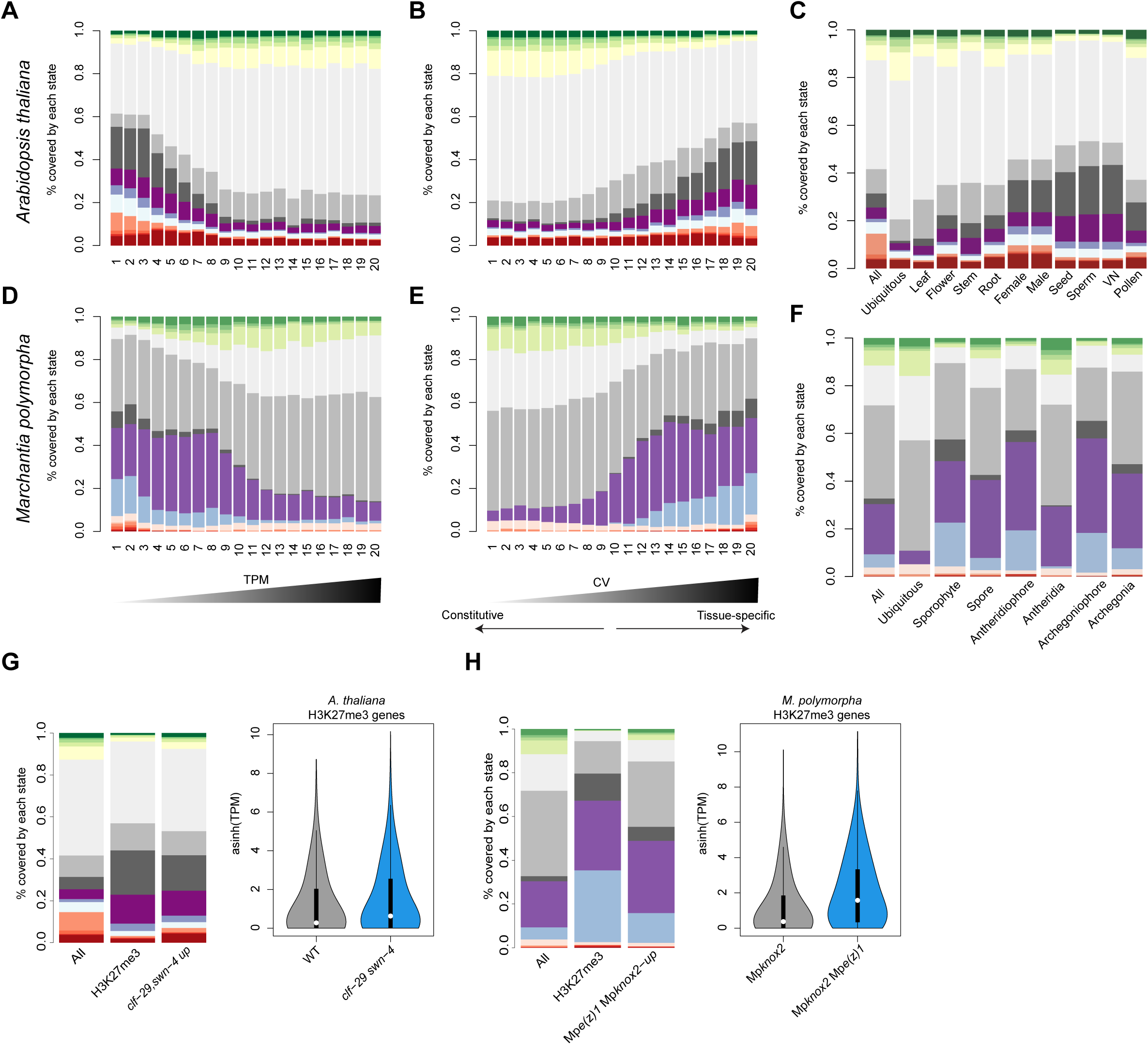
Association between chromatin states and transcription across promoters in *Arabidopsis* and *Marchantia*. **(A, D)** Bar plot of the proportion of chromatin states for 20 bins across promoters of genes ordered by transcripts per million (TPM) *Arabidopsis* leaves **(A)** and *Marchantia* thallus **(D)**. **(B, E)** Bar plot of the proportion of chromatin states across promoters for 20 bins of genes ordered by coefficient of variation (CV) across different tissues of *Arabidopsis* **(D)** or *Marchantia* **(E)**. **(C, F)** Bar plot the proportion of chromatin states across promoters for genes all genes or differentially expressed in specific tissues in *Arabidopsis* **(C)** and *Marchantia* **(F)**. **(G, H)** Bar plot of the proportion of chromatin states across promoters for genes covered by H3K27me3 or differentially expressed in PRC2 mutant in *Arabidopsis* **(G)** and *Marchantia* **(H)**. *clf/swn* double mutant for *Arabidopsis* and *Mpknox2 Mpe(z)1* double mutant for *Marchantia*, all genes are shown as a reference in both cases. In the right, asinh(TPM) of expression values for the same group of genes are shown for both mutant and the WT. The colour code for the diagram represents the chromatin states in the 15 states models.

To determine whether facultative chromatin states are associated with specific tissues, we examined genes with tissue-specific expression in both species. Contrasting with ubiquitously expressed genes, promoters of these genes exhibited a strong association with facultative chromatin states [Figures 3C and 3F], supporting the hypothesis that facultative heterochromatin states are associated with cell differentiation and developmental transitions, with a prevalence of H3K27me3 enriched states. The loss of this marks resulted in strong ectopic expression of the genes covered primarily by both types of facultative heterochromatin [Figures 3G and 3H].

Besides differences related to the organisation of chromatin states on expressed genes and the relative extension of facultative heterochromatin marked by H3K27me3 versus H2AUb, our analyses conclude a broad conservation of chromatin states composition, and their association with transcriptional activity of genes and silencing of transposable elements in *Marchantia* and *Arabidopsis*.

## Transcription Factors associate with specific chromatin states of *Arabidopsis*

The specific patterns of enrichment of histone modifications around the TSS both in expressed and repressed genes raised the possibility that specific transcription factors may preferentially associate with specific chromatin states. To investigate this hypothesis, we examined the genome-wide association patterns between transcription factors and chromatin states. To investigate TF binding preferences, we compiled a comprehensive dataset of ChIP-seq, DAP-seq, and position weight matrix (PWM)-based datasets from *Arabidopsis* and assigned TF-bound regions to chromatin states. ChIP-seq peaks showed a strong preference for intergenic states (especially I3 and I4) and E1 (associated with the TSS) and were depleted in constitutive heterochromatin and euchromatin states present on bodies of expressed genes but interestingly some TF enrichment was found in states of facultative heterochromatin [Figure 4A], to a large extent consistent with the accessibility constraints [Figure 1A]. Prediction of TF binding sites using DAP-seq, which does not depend on chromatin accessibility (42), exhibited a broader distribution across chromatin states with some preference for intergenic states [Figure 4B]. In contrast, PWM-based TF binding predictions, which only rely on DNA binding motifs, displayed a nearly uniform distribution [Figure 4C].

**Figure 4.**
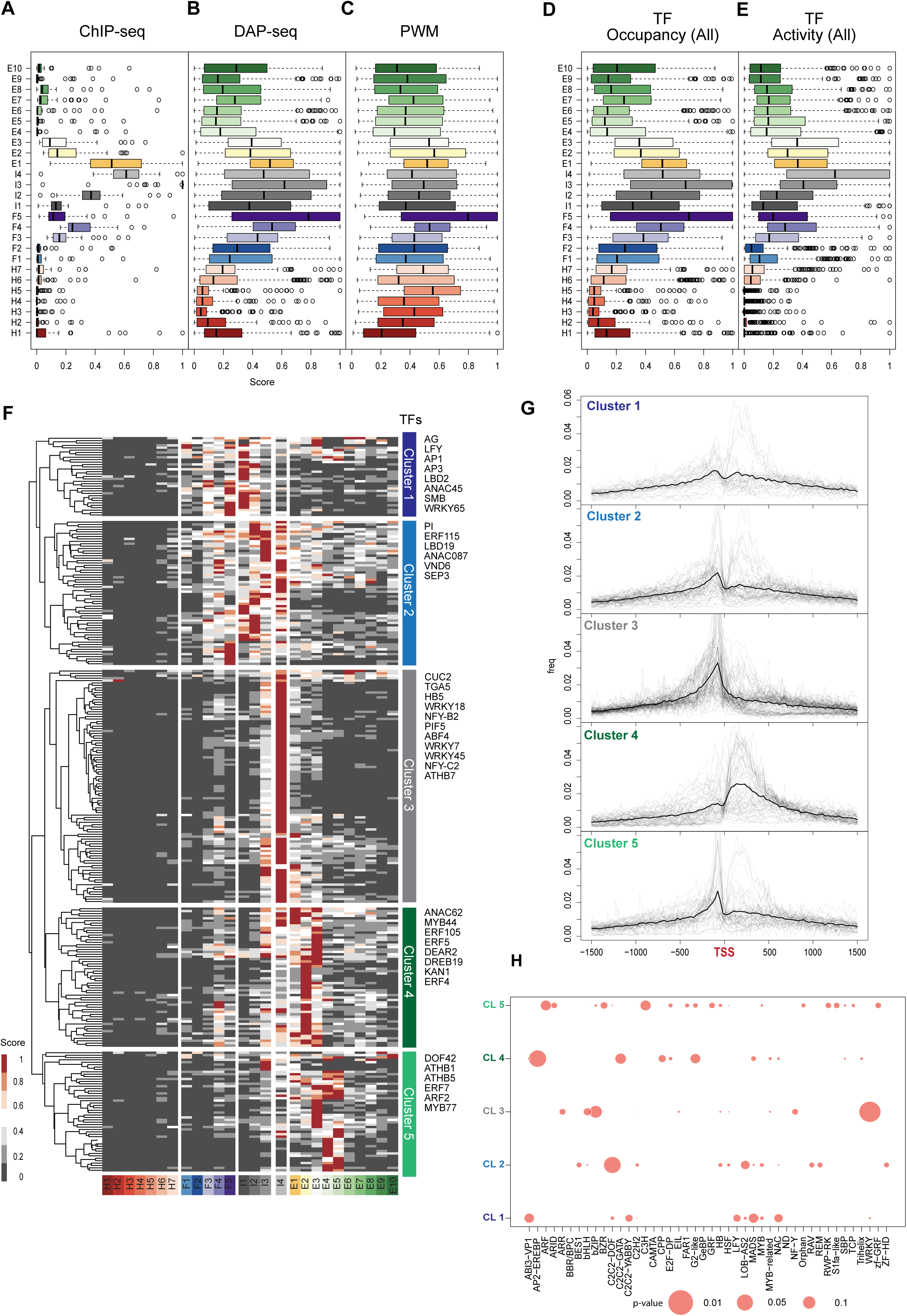
Transcription factors activity shows heterogenous chromatin state preferences in *Arabidopsis*. **(A-C)** Boxplot of TF occupancy scores based on ChiP-seq **(A)**, DAP-seq **(B)**, or position weight matrixes (PWM) motifs **(C)** in *Arabidopsis*. **(D, E)** Comparison of TF occupancy **(D)** and TF activity **(E)** scores across different chromatin states. The later scores are calculated as the statistical enrichment of TF binding over putative targets (co-expressed genes). **(F)** Heatmap and hierarchical clustering of TF activity scores for ChIP-seq and DAP-seq experiments across chromatin states. Colour scale is shown at the bottom. Scores are normalized to the maximum value of each TF. Low unnormalized scores were discarded. The colour code for the diagrams represents the chromatin states in the current 15 states models. **(G)** Histogram of TF binding distribution relative to the transcription starting site (TSS) for *Arabidopsis thaliana* TFs from different clusters defined in figure-H. **(H)** Bubble plot of Fisher’s Test of TF families across clusters defined in figure-H.

The observed differences between experimental approaches highlight the need for integrative analyses to fully resolve TF-chromatin state relationships in plants. TF occupancy combining data from ChIP-seq and DAP-seq was evaluated using an approach similar to that applied in human TF studies (5). To extend our analysis, we assessed TF activity, incorporating co-expression data as a proxy for functional interactions (55). We adapted a method to calculate chromatin state preference scores by evaluating the enrichment of TF binding to their putative targets, positively and negatively co-expressed genes (see Methods). Overall, low scores of TF occupancy and TF activity were broadly correlated with in heterochromatin states, and the highest scores were observed in chromatin accessible regions such as the intergenic states [Figures 4D and 4E], which is the main factor that differentiated ChiP-seq from DAP-seq and PWM [Figures 4A - 4E]. With this method to estimate TF activity, the scores of TF occupancy and activity converged. To look at different patterns of chromatin preferences among TFs, we kept ChIP-seq and DAP-seq data for ∼300 TFs in *Arabidopsis* (after filtering out TFs with low scores of occupancy and activity). Using hierarchical clustering, we identified distinct associations between TFs and chromatin states [Figure 4F] with distinct binding preferences relative to the TSS [Figure 4G].

Unlike *Arabidopsis*, *Marchantia* lacks a comprehensive dataset of ChIP-seq or DAP-seq experiments. However, previous studies have shown that TF DNA binding preferences are highly conserved across land plants (56), allowing to derive PWMs in *Marchantia* from PWMs from orthologous *Arabidopsis* proteins. Here, we combined recently published 24 DAP-seq experiments for *Marchantia* TFs with PWM from protein-binding microarrays (57) to serve as an alternative approach to infer TF binding. As with *Arabidopsis*, PWM-derived occupancy was evenly distributed across chromatin states [Supplementary Figure S2H]. However, when TF activity was estimated using co-expression, a striking shift emerged [Supplementary Figure S2I]. The most accessible chromatin states mI2 and mF2 were the most enriched, followed by mE2, mirroring the patterns observed in *Arabidopsis*. A clustering analysis revealed further preferential association of clusters of TFs with states enriched either before the TSS (I1 and I2) or after the TSS (F2 and E2) [Supplementary Figure S2J]. However, the scarcity of the data and the fact that we compared TF binding in diploid tissues of *Arabidopsis* with haploid tissues in *Marchantia* limited our analysis and further investigation is required to determine whether similar regulatory strategies are employed in the interplay between TFs and chromatin states to establish the degree of conservation of these interactions across land plants.

Remarkably in *Arabidopsis,* many TF families showed preferential association with one or two clusters [Figure 4H]. The largest group (Cluster 3, 32% of all TFs) was associated primarily with I4 and bound upstream of the TSS [Figure 4F and 4G]. This cluster comprised TFs from the WRKY, bHLH, bZIP, and HB families that bound to open chromatin of expressed genes (state I4) to regulate gene expression [Figure 4H and Supplementary Table 3], likely following the classic mode of RNA Polymerase II recruitment and stabilization of the pre-initiation complex (58). A GO term analysis indicated that TFs of cluster 3 primarily participated in biosynthetic processes, photosynthesis and circadian rhythm, and other housekeeping functions [Supplementary Table 7]. Other TFs associated with euchromatic chromatin states formed clusters 4 and 5. Notably, TF families present only in cluster 5 were involved in hormone signalling and flower development [Supplementary Table 9], including ARF (59). Most TFs from cluster 4 clearly bound after the TSS and several of these were associated with the control of the cell cycle [Supplementary Table 8], including E2FA, which is a typical marker of cell division. Clusters 4 and 5 also included the recently described GATC binding TFs (60), a large fraction of the AP2/EREBP family involved in stress response (61,62) and a large fraction of C2C2-GATA and G2-like TFs.

On the other hand, the TFs of Cluster 1 and 2 preferentially bound sites associated with facultative chromatin states [Figures 1A and 4F]. Cluster 2 contained TFs primarily controlling development, and more particularly root and vascular development [Supplementary Table 6]. Hence these TFs presumably control genes, which are not strongly active in leafy tissues of seedlings, in agreement with cluster 2 TFs occupying facultative heterochromatin states I1-I3 [Figure 4F]. Many of these TFs binding sites were located upstream of the TSS [Figure 4G] and are thus expected to bind to typical promoters. In contrast, cluster 1 TFs primarily associated with the first nucleosome downstream of the TSS [Figure 4G]. Most of these TFs controlled meristem and flower development and were significantly enriched in MADS-domain TFs, including APETALA1, PISTILLATA, APETALA3, SEPALATA3, and the pioneer factor LEAFY (LFY), which regulates flower development. Together with their target genes, these TFs are not expressed in seedlings where the chromatin states were described (22,63) [Figure 4F and Supplementary Table 5], which aligns with their binding to facultative heterochromatin. Overall, our results demonstrate that chromatin states can serve as a framework to classify TF-chromatin interactions. This analysis revealed distinct regulatory strategies among plant TFs involving proximal promoters in intergenic regions, the +1 nucleosomes, or cis elements downstream of the TSS most likely acting as enhancers or silencers.

## DISCUSSION

Here, we identified chromatin states in vegetative tissues of the liverwort *Marchantia* and found that their composition and association with transcription are strikingly conserved with the chromatin states of *Arabidopsis*. Overall, orthologous H2A variants and histone PTMs in *Marchantia* show the same associations and are equally important in defining chromatin states as has been shown in *Arabidopsis* (15). The variants H2A and H2A.X and the PTM H3K36me3 mark euchromatin; H3K27me3 and H2A.Z mark facultative heterochromatin; in constitutive heterochromatin, H3K9me1/2 associates with the variants H2A.W and H2A.M, which share C-terminal motifs enriched in lysine residues (20). Over protein-coding genes, chromatin states assemble in staggered arrays from the TSS to the TTS. Based on the integration of several datasets to map TF binding, we identified the preference of certain families and functional groups of TFs for specific chromatin states present either in the 5’UTR, the +1 nucleosome, or downstream of the TSS, suggesting a high complexity of cis elements controlling gene expression and the involvement of chromatin states in TF recruitment, either through direct recognition of chromatin marks or through co-regulatory mechanisms

The evolutionary distance between the two species and their distinct genome organization makes *Marchantia* an ideal model for exploring the conservation of chromatin states and highlighting the evolution of fundamental chromatin regulatory mechanisms. Overall, the organization of chromatin states in *Marchantia* confirms our previous evaluation of profiles of chromatin modifications and their conservation amongst bryophytes (50). Similarly, chromatin states described in *Arabidopsis* are representative of flowering plants (64,65). We observed species-specific heterochromatin organization, including the dispersed constitutive heterochromatin, the less complex and less abundant facultative heterochromatin in *Marchantia* marked by H3K27me3, a predominant share of regions with this mark also associated with TE in bryophytes (66). In *Marchantia*, euchromatin organization differs from *Arabidopsis*. The marks H3K4me1 and H3K36me3 reflecting transcriptional elongation and confined to the gene bodies in *Arabidopsis*, extend beyond the TTS in *Marchantia*, suggesting that signals for transcriptional termination differ between flowering plants and bryophytes. Overall, together with the similarities in the organization of histone modifications among bryophytes (50) and flowering plants (15,65) our data support the conservation of chromatin states and their relationship with gene expression in land plants. It is remarkable that this conservation is seen when comparing haploid gametophytic tissues of bryophytes and diploid sporophytic tissues of angiosperms, suggesting that chromatin organization in extant land plants reflects the organization of chromatin states in the haploid ancestors of land plants. The broad conservation of active chromatin states is also supported by a recent survey of chromatin states across eukaryotes while plants have evolved specific forms of repressive chromatin (67). Specifically, we show a broad independence between chromatin states governed by PRC1 and PRC2 in *Marchantia*, which contrasts with animals that connect the activities of the two modifiers through hierarchical recruitment models of “writing” and “reading” histone modifications H2AK121Ub and H3K27me3 (68,69). These differences support conclusions from phylogenetic analyses, which place the origins of the two independent types of Polycomb activities in the LECA, followed by divergence of their associations during evolution of different eukaryotic groups (70). Our results and the differences in the phenotypes of the mutants in PRC1 and PRC2 in *Arabidopsis* (71) further support the conclusion that PRC1 and PRC2 have remained largely independent from each other in plants.

In animals, distinct forms of association with chromatin distinguish functional types of TFs, including those associated with facultative heterochromatin (4,6). Comparable systematic studies were missing in plants. Here we integrate stable TF-chromatin interactions identified with ChIP-seq with transient, context dependent interactions and show that plant TFs associate with preferred chromatin states. Many TFs appear to follow a "canonical" mode of action, binding to open chromatin and regulating transcription by facilitating the assembly or stabilization of RNA polymerase II and the initiation of transcription [Supplemental Figure S5]. In *Arabidopsis*, this mechanism is predominant among TFs that regulate constitutive genes or modulate expression quantitatively. However, TF binding in *Arabidopsis* is not always associated with measurable transcriptional changes (24). Notably, open chromatin is potentially occupied by multiple TFs, suggesting extensive competition for binding. This could be due to open chromatin, which is associated with highly expressed genes and permissive for TF binding, generating highly occupied target regions (HOT) with redundant or passive activity (19).

Genes that require tight spatiotemporal regulation—remaining off in most tissues and activating only under specific developmental or environmental conditions—are more likely controlled by TFs capable of acting within facultative chromatin states. These TFs may interact with chromatin remodelers to induce local chromatin accessibility or directly access DNA protected by nucleosomes, resembling the pioneer TF paradigm described in animals (72,73). LFY functions as a pioneer TF by binding nucleosome-occupied sites at its targets APETALA1 and AGAMOUS, altering chromatin accessibility via SWI/SNF chromatin remodelers (20,74). MADS-domain TFs, bZIP TFs, LEAFY COTYLEDON1 (LEC1), TERMINAL FLOWER 1, FLOWERING LOCUS T, and MONOPTEROUS, have been proposed as candidate pioneer factors (75) and most belong to cluster 1 or 2, suggesting that these clusters define a new list of candidate pioneer factors.

Thus, TF preferences for specific chromatin states provide a sequence-independent axis for classifying plant TFs and inferring their regulatory modes. Generalizing this observation, our work captures the chromatin preference signature of pioneer TFs and expands the list of candidate pioneer TFs in plants.

## RESOURCE AVAILABILITY

Further information and requests for resources and reagents should be directed to and will be fulfilled by the lead contact, Frédéric Berger.

## DATA AND CODE AVAILABILITY

Datasets from the analyses are provided as supplementary data files. Pipeline and scripts used for data analysis are available on the following GitHub link: https://github.com/Gregor-Mendel-Institute/vikas_states_tf_study_2025.git. NGS data are deposited in GEO under accession numbers: GSE302232.

## Supporting information

supplementary tabbles

additional bed files

## ACKNOWLEDGMENTS

We thank the entire Berger group, for their insightful, considerate, and helpful discussions, as well as Matt Watson for editorial assistance during the preparation of the manuscript. We thank the Molecular Biology Service and Media Kitchen for a constant supply of plates, basic reagents, and cloning and sequencing services, as well as the Peptide Synthesis Service. Additionally, we thank the Vienna BioCenter Core Facilities, in particular the Next Generation Sequencing facility for their advice and swift handling of all our requests and helpful discussions. For open access purposes, the authors have applied a CC BY public copyright license to any author accepted manuscript version arising from this submission.

## AUTHOR CONTRIBUTIONS

Conceptualization: FB, and FR; Methodology: VS, FR, EA, and TH; Validation: VS, FR, and EA; Formal Analysis: VS, FR, and EA; Investigation: VS, FR, EA, and TH; Data Curation: VS, FR, EA, and TH; Writing – original draft: FB, VS, and FR; Visualization: ZHH, KS, and TW; Supervision: FB; Project Administration: FB, and FR; Funding Acquisition: FB, TH and FR.

