## supplementary figures for "Conservation of chromatin states and their association with transcription factors in land plants"

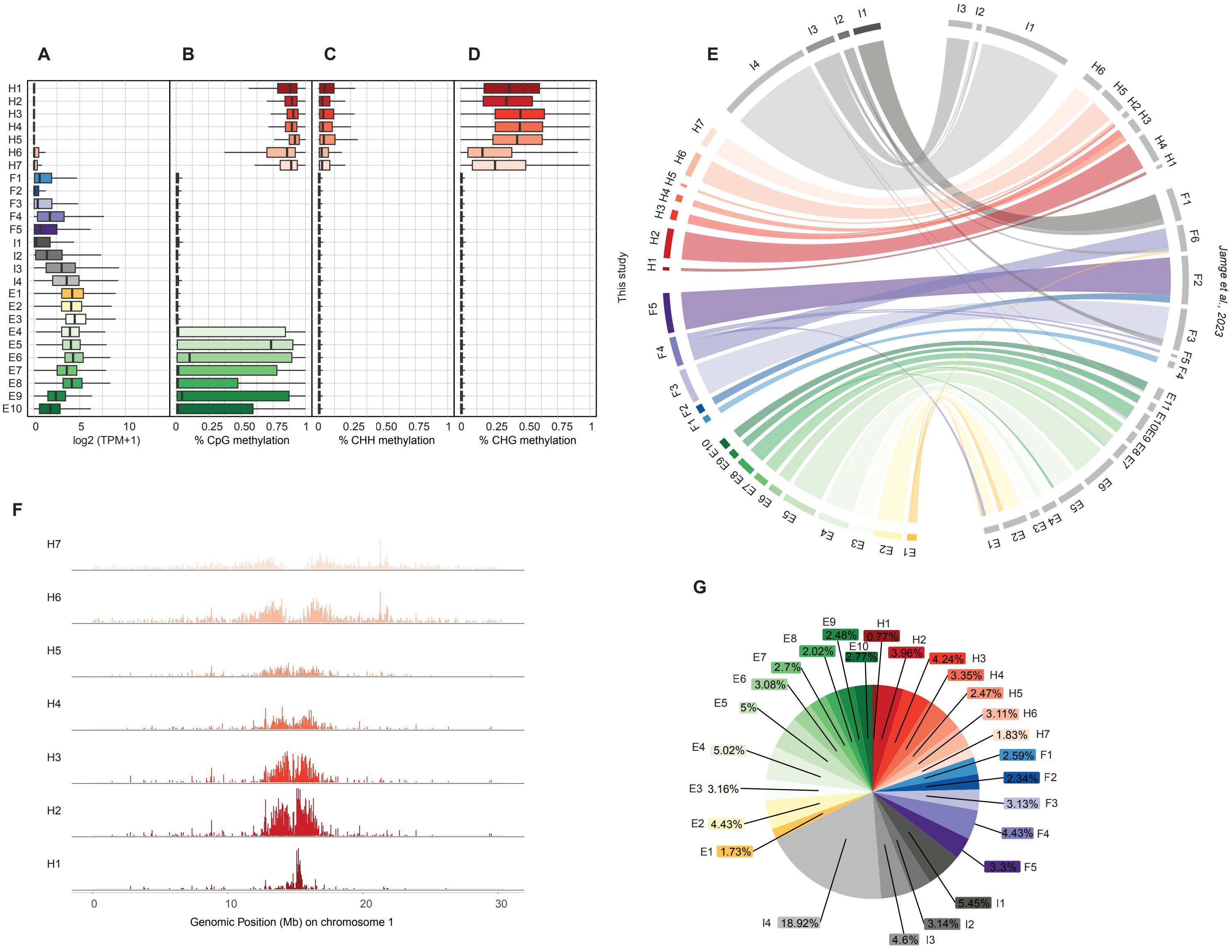

**Supplementary Figure S1**

**(A)** Box plot showing the expression of protein-coding genes overlapping with each chromatin state in Transcripts per Million (TPM). **(B-D)** Box plot showing levels of CpG **(B)**, CHH **(C)**, and CHG **(D)** methylation for all chromatin states. **(E)** Flow diagram showing the overlap (in bp) between the chromatin states defined in this study with states from Jamge et al., 2023. The colour code for the flow diagram represents the chromatin states in the current model. **(F)** Genomic distribution of heterochromatic states (H1–H7) [bottom to top] on Chr1 of Arabidopsis thaliana. **(G)** Pie chart showing the percentage of the genome covered by each state.

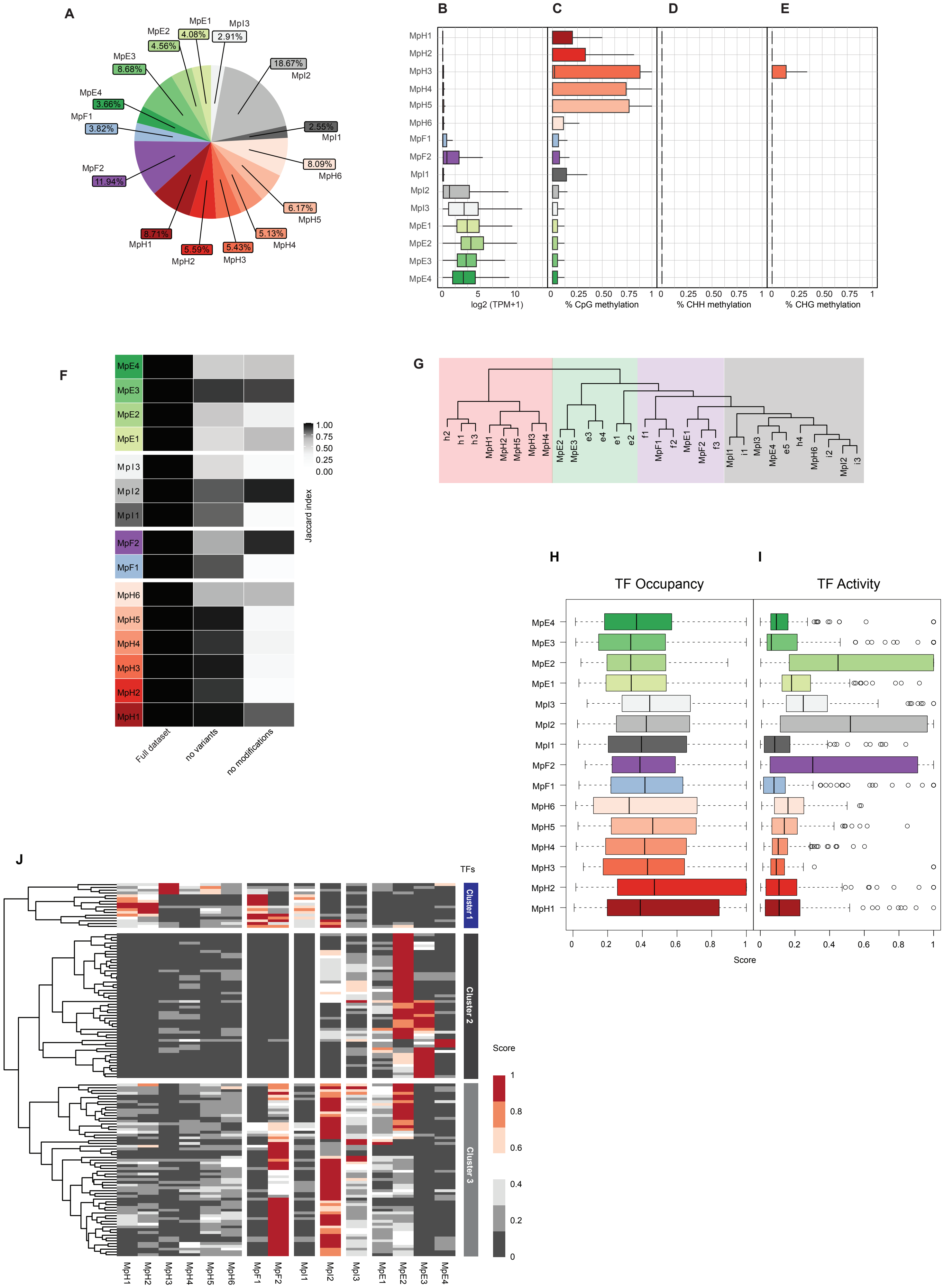


Supplementary Figure-S2

**Supplementary Figure S2**

**(A)** Pie chart showing the percentage of the genome covered by each state of *Marchantia polymorpha*.

**(B)** Box plot showing the expression of protein-coding genes overlapping with each chromatin state of *Marchantia polymorpha* in Transcripts per Million (TPM).

**(C-E)** Box plot showing levels of CpG **(C)**, CHH **(D)**, and CHG **(E)** methylation for all chromatin states of *Marchantia polymorpha*.

**(F)** Heatmap showing the Jaccard similarity index between the states generated using the whole model and states using a subset of marks, i.e., excluding a set of marks and variants as indicated on the x-axis.

**(G)** Hierarchical clustering of the emission probabilities of chromatin states of *Marchantia* and a down-sampled 15-states model of *Arabidopsis* representing the conservation of chromatin state definition in the two distantly related species.

**(H, I)** Boxplot of TF occupancy **(H)** and TF activity **(I)** scores across different chromatin states. The later scores are calculated as the statistical enrichment of TF binding over putative targets (co-expressed genes).

**(J)** Heatmap and hierarchical clustering of TF activity scores for PWM across chromatin states. Colour scale is shown at the bottom. Scores are normalized to the maximum value of each TF. Low unnormalized scores were discarded. The colour code for the diagrams represents the chromatin states in the current 15 states models.

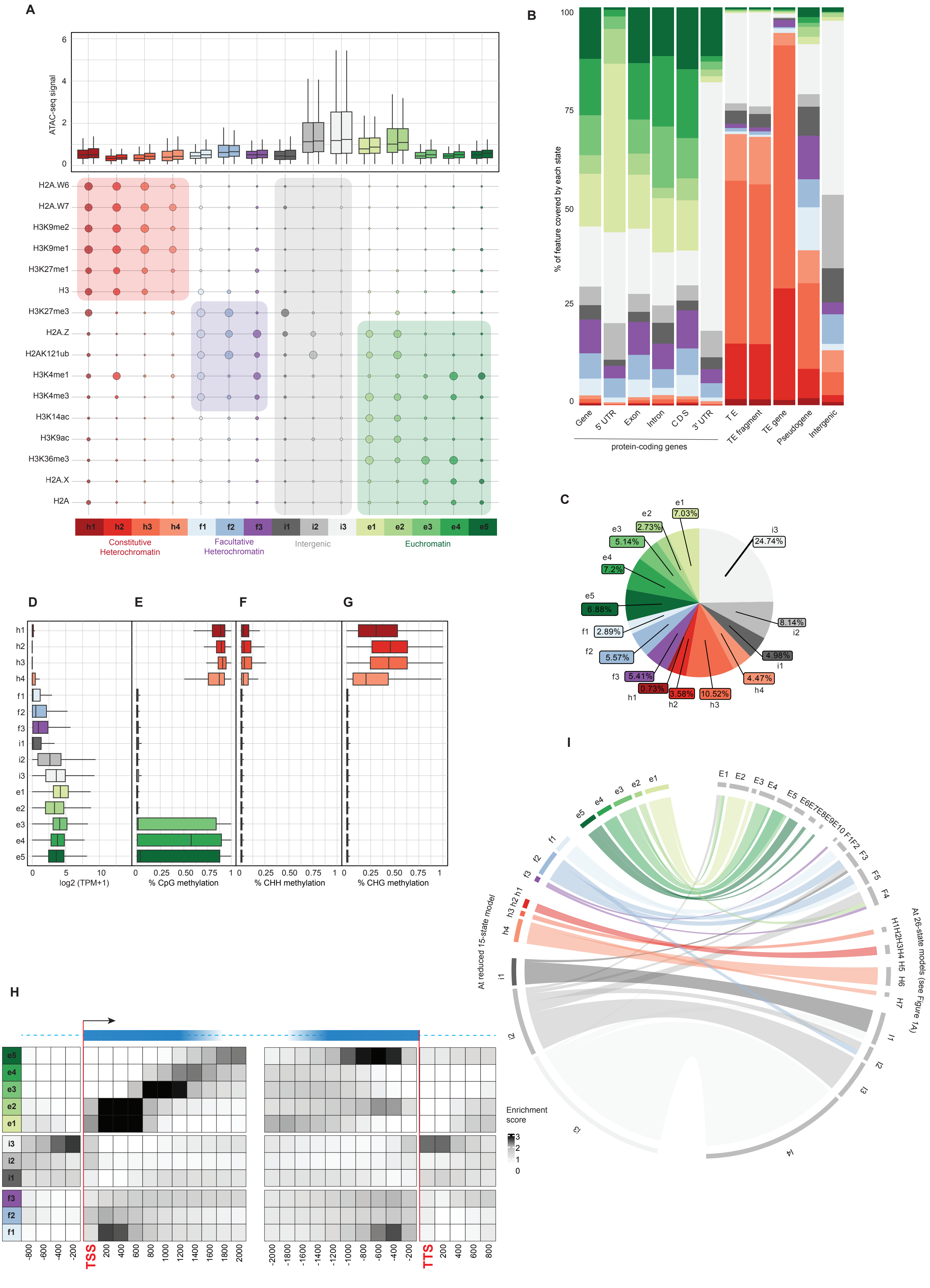


Supplementary Figure-S3

**Supplementary Figure S3**

- **(A)** Bubble plot showing the emission probabilities for histone modifications/variants for a down-sampled 15-state model of *Arabidopsis thaliana*. The size of the bubble represents the emission probability ranging from 0 to 1. Coloured rectangles demarcate the classification of states into major domains of chromatin. The box plot on top shows the average ATAC-seq signal for each state representing chromatin accessibility. The two boxes per state are two replicates of the ATAC-seq experiment.
- **(B)** Stacked bar plot showing the overlap between annotated genomic features and the chromatin states in the 15-state model of *Arabidopsis thaliana*.
- **(C)** Pie chart showing the percentage of the genome covered by each state of the 15-state model of *Arabidopsis thaliana*.
- **(D)** Box plot showing the expression of protein-coding genes overlapping with each chromatin state of the 15-state model of *Arabidopsis thaliana* in Transcripts per Million (TPM).

- **(E-G)** Box plot showing levels of CpG, CHH, and CHG methylation respectively, for all chromatin states of the 15-state model of *Arabidopsis thaliana*.

- **(A)** Neighbourhood Enrichment analysis of chromatin states of *Arabidopsis* 15-state model representing the fold enrichment for each state at fixed positions relative to the anchor position (TSS and TTS).

- **(B)** Flow diagram showing the overlap (in bp) between the chromatin states of extensive 26-state model and the 15-state model *Arabidopsis thaliana*. The colour code for the flow diagram represents the colours for the states of the 15-state model.

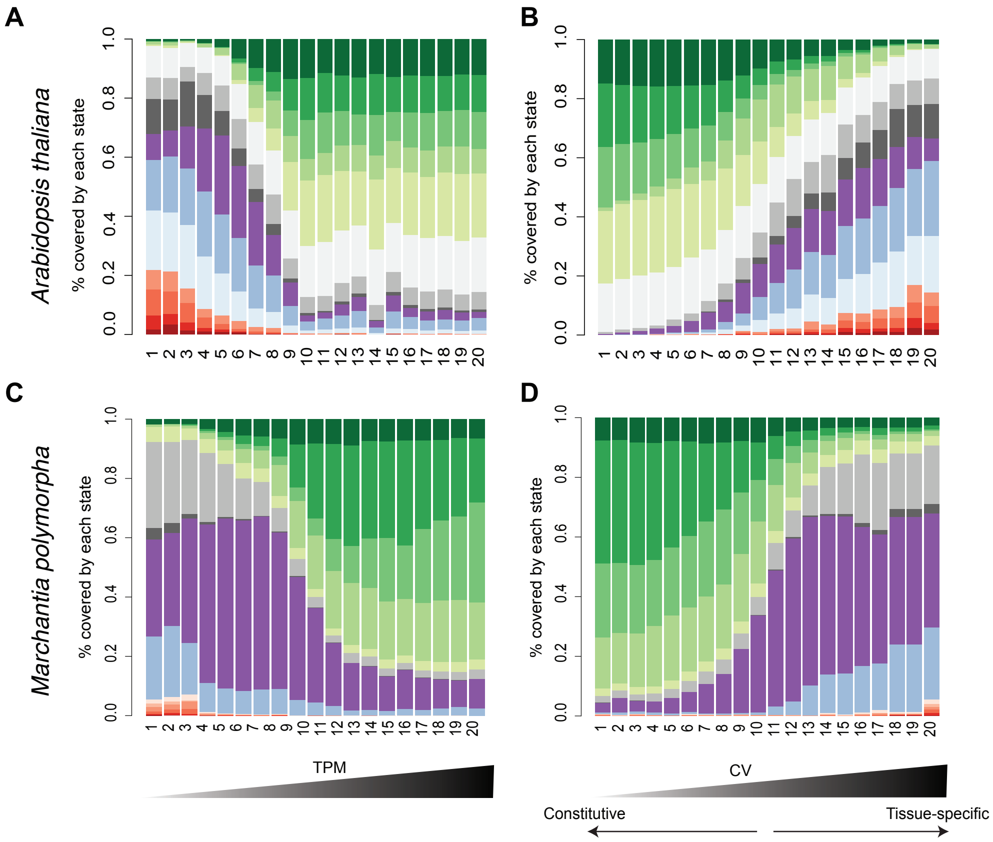


**Supplementary Figure-S4**

**(A-D)** Bar plot of the proportion of chromatin states for 20 bins across coding regions of genes ordered by transcripts per million (TPM) Arabidopsis leaves **(A, B)** and Marchantia thallus **(C, D)**.

A

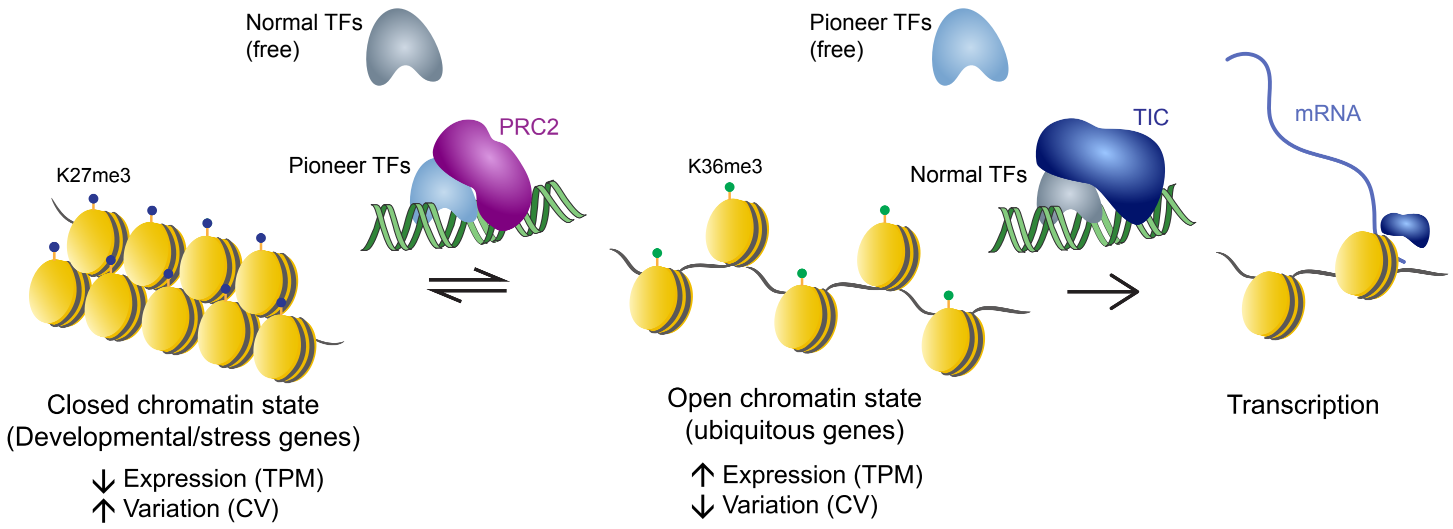


**Supplemental Figure-S5**

**(A)** Model of transcription factor interplay with facultative and open chromatin to control transcription rates.
